# A theory of neocortical seizure spread: Insights from statistical physics

**DOI:** 10.1101/691576

**Authors:** Cole A. Giller

**Author notes:** Corresponding author (CG).

## Abstract

The conception of seizures as abnormal synchronies of large neuronal populations has been confirmed by numerous electrophysiological studies, including recent imaging of travelling seizure waves across the neocortex. This traditional viewpoint has been challenged by the finding that during some seizures, neurons with high firing rates are remarkably rare and sparsely distributed into clusters. Reconciliation of these seemingly contradictory descriptions has attracted much attention, raising questions such as how (or if) macroscopic seizure waves arise from these microscope neuronal clusters, and more generally, how other features of macroscopic, clinical seizures arise from microscopic dynamics. Answers to these questions are crucial to the understanding of epilepsy, and could guide development of drugs and other interventions that act at the microscopic level to effect macroscopic improvement.

Relationships between microscopic and macroscopic processes are addressed by the field of statistical physics, offering explanations for how macroscopic quantities such as pressure and temperature arise from microscopic interactions between molecules. Here we hypothesize that these methods could also provide insight between the macroscopic and microscopic dynamics of seizure behavior. We constructed a model of the neocortex composed of small domains, each representing a cluster of neurons. Models with and without refractory periods were studied. Allowing seizures to spread among the clusters in a probabilistic fashion produced a “cellular automaton” amenable to the methods of statistical physics. We thereby showed that the model harbors a continuous phase transition allowing possible explanations for the emergence of seizure waves from microscopic neuronal clusters, and for a surprisingly wide variety of seizure properties. Moreover, the model is easy to use because it requires only a small number of intuitively understood rules and is computationally efficient. We hope that these insights from statistical physics will contribute to the understanding of epilepsy and to the identification of new therapeutic measures.

**Author summary:** Epilepsy is a common neurological disease characterized by devastating, unpredictable seizures. Extensive research is aimed at improving the treatment of epilepsy through better understanding of how seizures start and spread, but basic questions remain unanswered. Do seizures start as waves of overactive neuronal activity, or as small clusters of activity as suggested by recent data? How do clinical properties of seizures emerge from interactions between small groups of neurons? And would understanding this emergence lead to better treatment?

We address these questions with a mathematical model of seizure spread, using methods of physics designed to explain how quantities such as pressure and temperature emerge from interactions between molecules. The model produced small clusters of activity as observed in recent data, and the methods allowed us to show how these clusters react to increases in neuronal excitation to produce seizure waves and other clinical seizure behavior. The model thus provided possible answers to the questions above, based on new insights from the field of physics. If the model indeed represents a common pathway evoked by many pathological changes, it may inform the development of therapeutic measures such as antiepileptic drugs that act at the microscopic level to improve macroscopic behavior.

## Introduction

Epilepsy is a disabling disease in which the normal balance of synchrony between neural networks within the brain is replaced by an intense, often widespread synchrony that overwhelms normal function and produces the clinical symptoms of seizures. Treatment includes the use of drugs acting at a microscopic level to curtail the macroscopic seizures, and the use of surgery to resect seizure ‘foci’ or disrupt macroscopic networks responsible for seizure spread. Much work has therefore focused on how clinical, macroscopic seizures arise from interactions at the microscopic level between neurons, both to explicate the pathophysiology of epilepsy and to inform the design of drugs or other interventions that exert effects on microscopic neuronal dynamics.

A traditional approach to these issues has been to view seizures as hypersynchronous discharges of large neuronal populations, organized into waves and spirals similar to those observed *in vivo* and produced by computational models [1–6]. Challenging this approach, however, are studies showing that neurons with high firing rates during seizures are both rare and sparsely distributed [7–10]. Moreover, recent human data show that active neurons are organized into small, discrete domains with diameters similar to those of cortical columns, and that harbor brief bursts of ‘microseizures’ [11–18]. Furthermore, it has been shown that the neurons giving rise to markers of seizure activity such as high frequency oscillations and interictal spikes are organized similarly [19–21]. Hence there is credible but puzzling evidence that seizure activity manifests both as small clusters of hyperactive neurons and as large areas of continuous waves. The seeming discrepancy between these views emphasizes the importance of understanding the relationship between macroscopic and microscopic seizure activity, and in particular, understanding how macroscopic seizure waves arise from microscopic dynamics.

In order to understand these relationships, some have hypothesized that microseizures form ‘small clusters of pathologically connected neurons’, which coalesce to produce macroscopic, clinical seizures [22–26]. There is also evidence that the transition from microscopic dynamics to macroscopic waves occurs when a small ‘ictal core’ of wave activity strongly inhibits a surrounding area of sparsely distributed active neurons, and that EEG waves are associated with this inhibition [11, 27]. Others suggest that the transition occurs through increases in extracellular potassium or increases in excitability [28, 29]. But it is still unclear how microscopic dynamics might produce clinical events such as sudden seizure cessation, preictal desynchronization, sensitivity to physiological parameters or the ways in which seizure foci affect seizure behavior.

Understanding dynamic processes at different spatial scales is amenable to mathematical modeling, and many such models have been used to explain the emergence of macroscopic seizure behavior from microscopic interactions. For example, several models characterize seizures by critical behavior, so that events such as seizure cessation can be viewed as ‘tipping points’ [30–33]. It has been difficult, however, to find a model that is both intuitive and capable of capturing more than a few of the key features of seizure behavior. Some models lack spatial relationships [34], some require interactions between every pair of cortical columns [35], and some lack clusters or spirals, or require many differential equations and many assumptions of parameter values [5, 6, 26, 36–40]. Other models address large-scale connectivity but depend on variables without obvious intuitive biological meaning [31, 41, 42].

Our goal in this work is to construct a model that requires only a small number of intuitively interpretable variables, but that nevertheless captures many of the features of seizure behavior mentioned above. We start with the observation that the field of statistical physics explains how macroscopic quantities such as pressure and temperature might arise from microscopic interactions between innumerable molecules [43–45], and we then hypothesize that in a similar fashion, these methods could explain the emergence of macroscopic seizure behavior from the interactions between large numbers of microscopic neuronal domains.

Our model consists of a two-dimensional grid of ‘domains’ of size approximate to that of cortical columns, each capable of harboring a microseizure. Allowing microseizures to spread probabilistically between adjacent domains renders this model into a cellular automaton that can be studied with tools of statistical physics, and that has been helpful in understanding how the macroscopic properties of crystals and magnetic domains arise from microscopic dynamics [46, 47]. We show that our model harbors a phase transition between the seizure and non-seizure states, much as some substances transform from liquid to solid form, that can be studied with methods borrowed from statistical physics. We then show that with the addition of refractory periods, the model offer explanations of how macroscopic seizures might arise from microscopic dynamics of domains, as well as a surprisingly wide variety of well-known seizure behaviors. These include the presence of enlarging clusters of microseizures, the common observation of sudden seizure onset and cessation, the presence of seizure waves, a high sensitivity to small changes in physiologic variables, the frequently observed prolonged and complex responses to transient stimulation, preictal desynchronization, reentrant loops, and the puzzling, paradoxical worsening of seizures that can occur with increased inhibition. Although we do not provide new experimental data, we believe that the sheer number of seizure properties that can be explained with the model supports the hypothesis that methods of statistical physics can provide valuable insights into seizure behavior and may suggest new approaches to seizure treatment.

## Results

### Description of Model

The model consists of a 2-dimensional rectangular patch of ‘neocortex’ composed of domains with diameters approximating those of cortical columns, arranged in a hexagonal grid based on estimates of column geometry [48, 49]. In simulations, the grids ranged in size from 200×200 to 1000×1000. A domain interacts only with its adjacent neighbors, and is *active* if it harbors a microseizure. Active domains appear as red in our simulations, and non-active domains as black. Domains on the patch boundary are held inactive to simulate a surrounding normal zone. Because a regular hexagon is wider than it is high, the patch appears in the simulations as a rectangle with a width:height ratio of 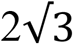.

Time advances in steps taken for a seizure to travel between two adjacent domains. At each step, a domain can enter the seizure state spontaneously with a probability of pr (“p random”), by remaining in the seizure state from one step to the next with a probability of ps (“p stable”), or by spread from an adjacent active domain with a probability of p (Fig 1).

**Fig 1.**
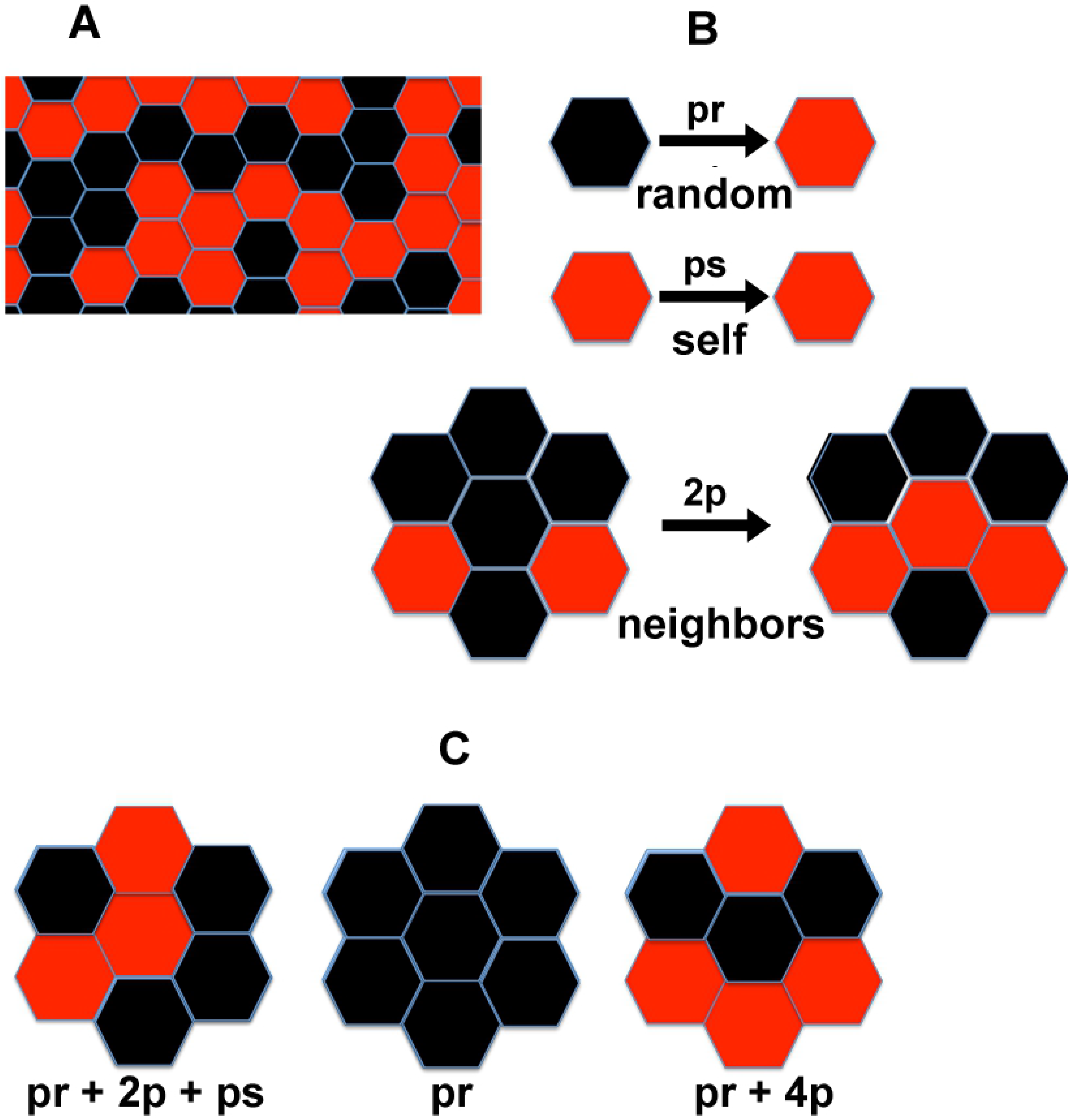
Local Probabilistic Interactions Between Domains. (A) Portion of grid showing activated domains in red and non-activated domains in black. (B) pr is the probability of a domain spontaneously activating; ps is the probability of an activated domain staying active at the next time step; kp is the probability of a domain activating due to its neighbors, where k is the number of activated neighbors. (C) Examples of probabilities of the central domain being active at the next time step.

The total probability of activation for each domain at each time step is additive, i.e., is pr + ps + kp whenever this quantity is ≤ 1. The probability ps is set to 0 if the domain was not active in the prior time step, and k is the number of active neighboring domains (Fig 1B). If pr + ps + kp exceeds 1, we define the transition probability to be 1.

These systems are *probabilistic cellular automata* [47], consisting of a lattice of nodes (domains) in different states (active vs. non-active), evolving in time through transition rules (activation with probability of pr + ps + kp). Cellular automata produce complex behavior from simple transition rules, allowing simulation of otherwise intractable systems [50].

### Percentage of Active Domains

We write m(t) for the percentage of active domains at time t, and m for the asymptotic value of m(t) as t becomes large. A model is called *saturating* if pr + ps + 6p ≥ 1, and non-saturating if pr + ps + 6p < 1. We show in the Methods section that all domains eventually activate for saturating models in which pr is non-zero, and that for non-saturating models, m is given by

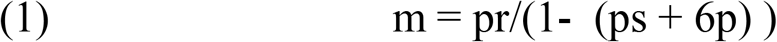

Closed form results such as Equation 1 for cellular automata are rare [51], but our simulations confirm this equation for both non-saturating and saturating models. For non-saturating models with pr = 0, no activation occurs and m = 0 as predicted. For non-zero pr, activation began spontaneously, and m(t) increases to oscillate around its asymptotic value m as predicted by Equation 1 (Fig 2). The oscillations are due to the probabilistic nature of the model and the effects of a finite grid size.

**Fig 2.**
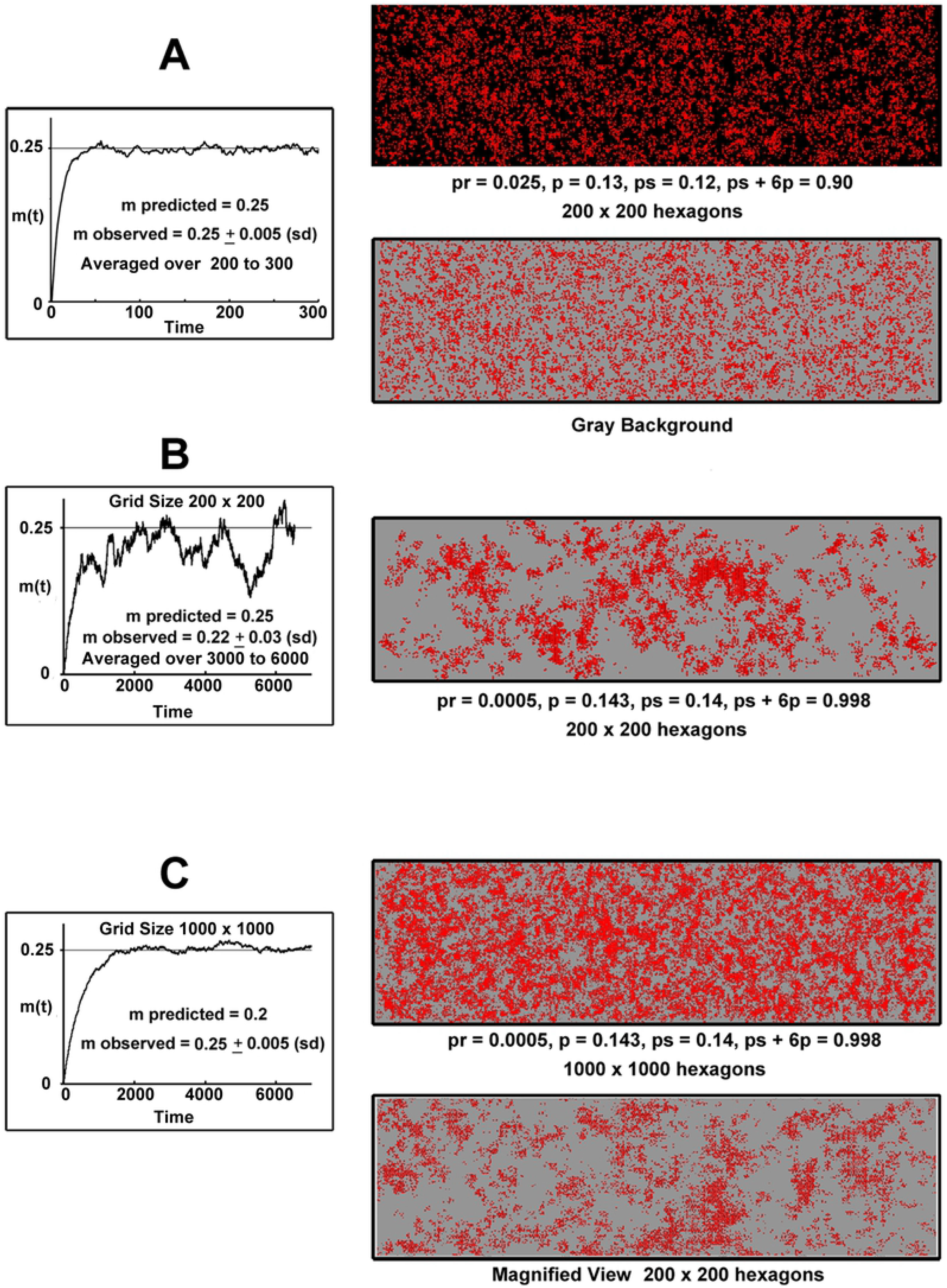
Effects of Excitability and Grid Size on Predicted and Observed Percentage of Active Domains. (A) Simulation in moderately excited field (ps + 6p = 0.90). The value of m predicted by Equation 1 agrees with the values averaged from time step 200 to 300. The time taken to reach equilibrium is relatively small. Gray background is used for clarity. (B) Simulation with greater excitability, but same value of m. Clusters of activated domains are larger, as expected for higher excitability. Variation in m averaged over time step 3000 to 6000 is greater due to small grid size (200 × 200 hexagons). (C) Same parameters as in (B), but the grid size of 1000 × 1000 is larger. The variation in m is smaller, and clusters remain large in total and magnified view. Note the longer time taken to reach equilibrium due to slowing down effects. Data are represented as mean ± SD.

### Phase Transitions and Criticality in Non-saturating Models

In contrast to sudden first order transitions between model states, *continuous* phase transitions occur gradually and have hallmark properties of sensitivity to small changes, ‘slowing down’ of dynamics, long-range correlations of domain states, enlarging clusters of activity, and complex responses to stimuli [30, 33, 43, 45, 52, 53]. In this section we show that our model has these properties, providing evidence for the existence of a continuous phase, and show how these properties can be translated into statements about seizure behavior. The transition occurs when c = ps + 6p approaches a *critical point* C_0_ slightly less than 1 − pr. We will say that the model is ‘near criticality’ when c is close to C_0_, which is itself close to 1 − pr. Critical behavior thus arises from increases in excitability due to increases in ps or p, but not from increases in pr.

#### Sensitivity to small changes

Phase diagrams derived from Equation 1 reveal that when the system is near criticality, m is highly sensitive to c and approaches 1 (Fig 3A). Moreover, for a fixed value of c close to 1 − pr, m is also sensitive to changes in pr. This can also be seen from the derivatives of m with respect to pr, p and ps, which are high when close to criticality (i.e., when 1 − c is close to pr). This sensitivity is the first hallmark of a phase transition. In the language of statistical mechanics, m is the *order parameter* of the system, the transition to global activation (m = 1) occurs as a continuous phase transition is approached, which in turn occurs when pr is small and c approaches 1 − pr.

**Fig 3.**
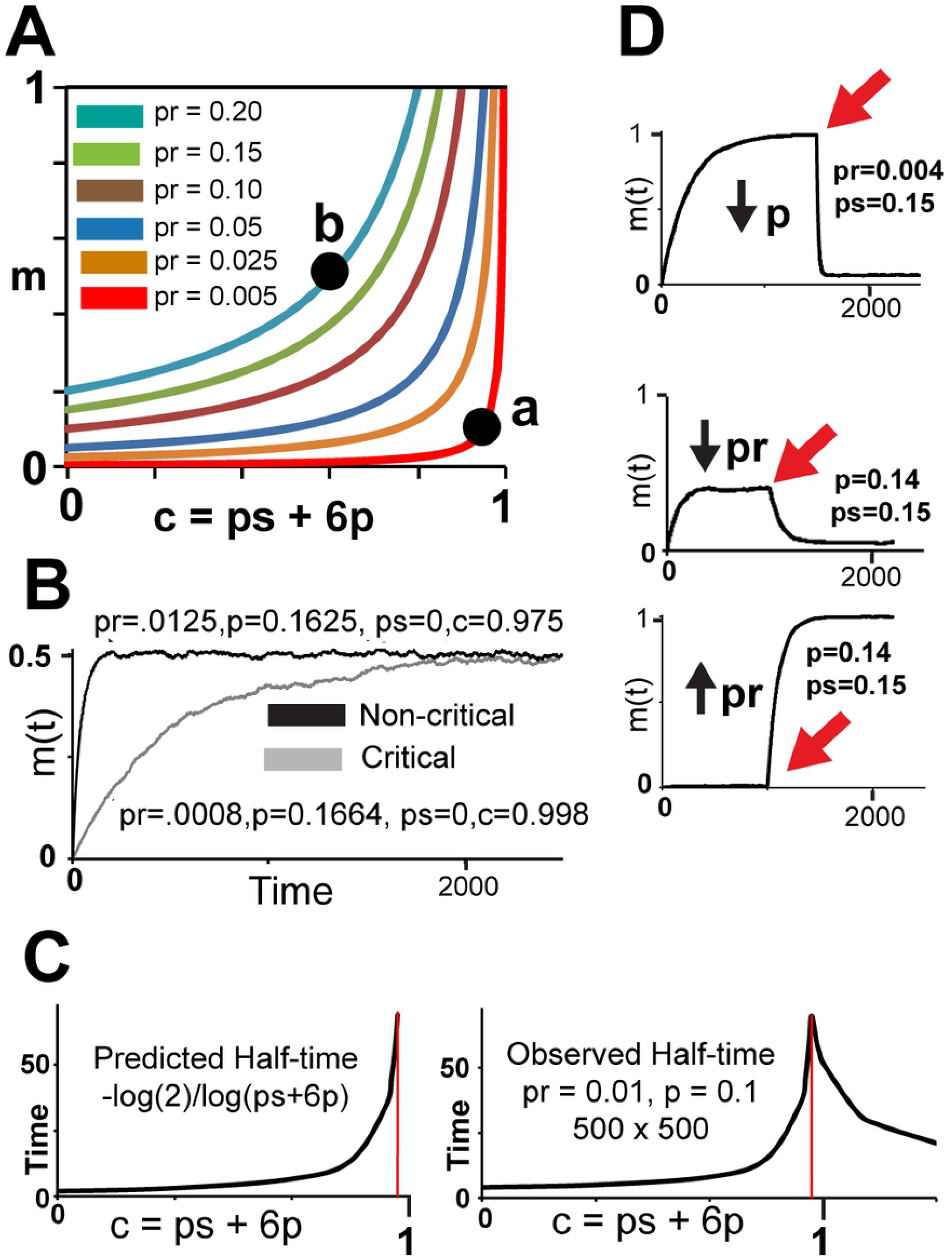
Effects of Criticality: Phase Diagram, ‘Slowing Down’, Sensitivity to Parameters. (A) Phase diagram showing rapid increase of m as c approaches 1 − pr. (B) m(t) of the system close to criticality takes longer to reach equilibrium than the non-critical system due to slowing down. Both systems have the same m from Equation 1 and the simulations (C) Predicted half-time vs. c (top) compared to simulations as ps increases with pr = 0.01 and p = 0.1 (bottom). ‘Slowing down’ occurs when c approaches 1 − pr from either direction. (D) m(t) vs. time for excitable systems. Top graph shows that a system in which a small decrease of p from 0.141 to quickly reduces m(t) from 1.0 to 0.06. Middle graph shows a system in which a small decrease of pr from 0.004 to 0.0005 quickly reduces m from 0.4 to 0.05. Bottom graph shows a system in which a small increase in pr from 0.0001 to 0.01 increases m(t) from 0.01 to 1.0.

#### ‘Slowing Down’ of dynamics

System dynamics ‘slows down’ near a continuous phase transition [33, 52], as observed during transition to neuronal spiking and the onset, spread and termination of seizures [24, 31, 32, 54, 55]. Building on previous reports [47, 56], we show (see Methods) that a non-saturating model ‘slows down’, i.e., that the time required for m(t) to reach its predicted value m grows exponentially near criticality, as in Fig 3B. Simulations also showed that the time taken to activate all domains rapidly decreases as c increases beyond 1 − pr, so that ‘slowing down’ occurs as c approaches 1 − pr from either direction. (Fig 3C). The ‘slowing down’ of our model is a second hallmark of a phase transition.

#### Long-range correlations of domain states

Fluctuations of domain states are correlated due to local interactions, but correlations decrease exponentially with distance. The decay constant is the *correlation length,* and grows without limit as criticality is approached [43, 45]. Cortical patches are finite, and so the peaks in the correlation graphs are also finite. To our knowledge, the correlation length of our model cannot be directly calculated, but we computed correlation lengths for several examples following Nagy et al. [57]. The finding of peaks in the graph of correlation length near criticality (Fig 4C) is a fourth hallmark of a phase transition.

**Fig 4.**
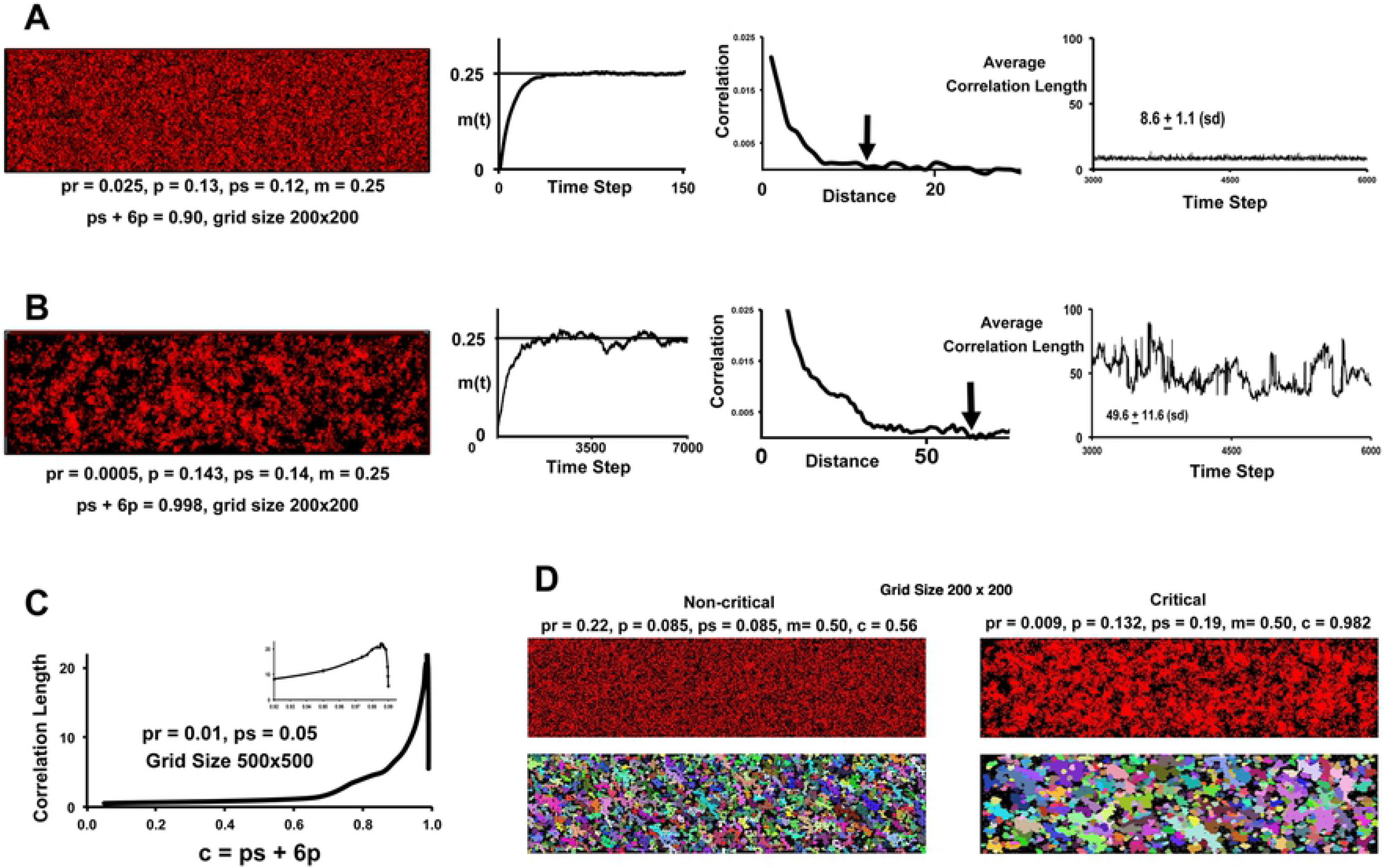
Correlation Length and Cluster Size (A) Calculation of correlation length for simulation with moderate excitability (c = 0.90). The graph on the left shows a fast rise of m(t) to its equilibrium value of 0.25, and the grid image was obtained after that time step. The average correlations of each domain with each of its four neighbors at successively larger distances were calculated, and these correlations plotted against distance (middle graph). The distance at which the correlation was first less than 0.001 was taken as the correlation length for the domain in question (arrow). Averaging over all domains within a central block, repeating the process for each time step (graph on right), and averaging over the last 2000 time steps produced the correlation length for the simulation (see Methods). As expected for moderate excitability, the correlation length in this example was low (8.6 ± 1.1 sd). (B) Same calculation for simulation with high excitability (c = 0.998). The correlation length is higher (49.6 ± 11.6), leading to the formation of clusters seen in the grid. (C) Correlation lengths for systems of increasing excitability (in this case, pr and ps were held at 0.01 and 0.05, respectively, and p gradually increased). The peak near 1 − pr as c increases is strong evidence of a continuous phase transition. The sharp decrease after the peak is due to the use of a finite grid. Inset shows details at peak, consistent with a phase transition near c = 0.99. (D) Simulations comparing a system far from criticality to a system near criticality. The mean cluster size is larger for the system near criticality. See text and Methods.

#### Enlarging clusters of activity

Increasing the correlation length also increases correlations between large groups of domains, resulting in the formation of clusters of activity that enlarge near criticality [45]. To illustrate, we compared 10 simulations of two systems with the same m in which one system was near criticality (Fig 4D). The number and size of clusters were counted using a density-based cluster algorithm (see Methods). The clusters of the non-critical system were significantly more numerous (1246.9 ± 20.4 vs. 683 ± 21.2, p < 0.001, mean ± sd) and smaller (13.5 ± 0.28 vs. 23.2 ± 1.1, p < 0.001) than those of the critical system. This fifth hallmark of phase transition also provides an explanation for the observed clusters of hyperactive neurons.

#### Complex responses to stimuli

For non-saturating models near criticality, ‘slowing down’ prolongs the response to a stimulus, and in some cases, thousands of time steps are required to approach m (Fig 3B and Fig 5). This suggests that seizures in the setting of low excitability may paradoxically achieve their maximum effect much sooner than when excitability is high, i.e., a long ‘buildup’ of seizure activity may have more severe clinical effects than a rapid onset of activity. For saturating models near criticality, ‘slowing down’ led to a slow onset of global activation. These complex responses to stimuli constitute a third hallmark of a phase transition.

**Fig 5.**
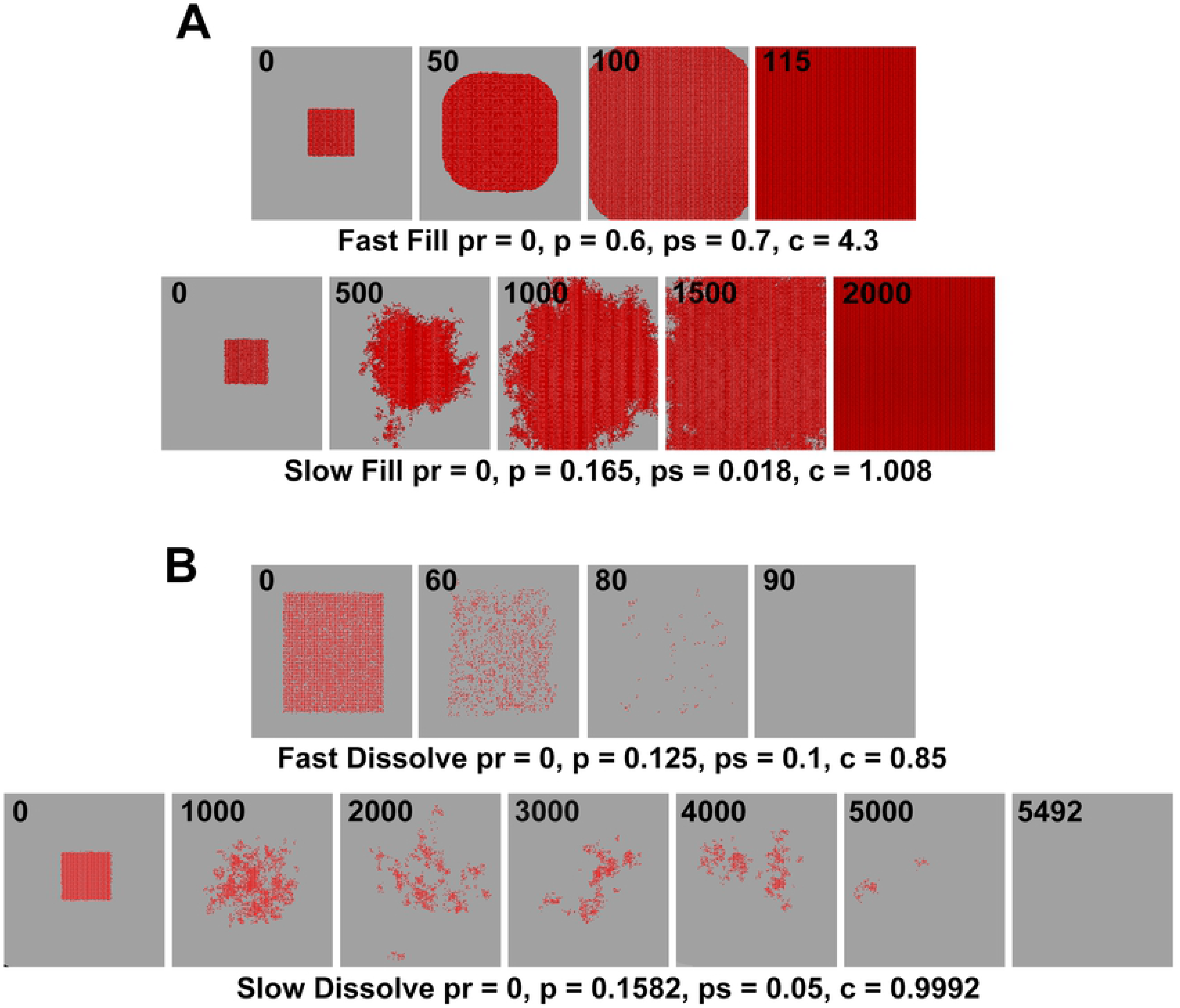
Effects of Criticality on Time Taken to Reach Equilibrium. (A) Response to transient focus when system is saturating. Both systems undergo global activation, but the system shown in the top row is far from criticality and fills at a much slower rate than the system near criticality on the bottom. (B) Response to transient focus when the system is non-saturating. In both cases all domains deactivate because pr = 0, but at a slower rate when the system is near criticality (bottom row). Effects of a focus can be complex near criticality, even when the asymptotic result is known.

In addition to these hallmarks, it is known that the time history of a probabilistic cellular automaton is equivalent to a 3-dimensional Ising model made of copies of the 2-dimensional automaton, ‘stacked’ along the time dimension, having continuous phase transitions that can be explicitly calculated [46, 47, 58]. Doing so for our model by following the appendix of [47] suggests a critical point near 1 − pr, in agreement with our other estimates.

These properties of non-saturating models thus support the presence of a continuous phase transition, occurring as ps or p increase enough to push c = ps + 6p close to the critical point of the system. It is important to note that the excitability of our model can increase in two distinct ways. The first occurs through increases in ps or p, i.e., by increases in the rate of seizure spread or persistence. This produces an increase in the percentage m of active domains according to Equation 1, moves the system towards criticality, and confers the hallmark properties of criticality as discussed above. The second way in which excitability can increase is through increases in pr, i.e., increases in the rate of spontaneous domain activation. This produces an increase in m according to Equation 1, but does not move the system towards criticality or induce properties of a continuous phase transition. These considerations may explain how two seizure types resulting in the same severity of symptoms (i.e., having the same values of m) may have very different sensitivities, timings and patterns of activation depending upon which type of excitability (increases in pr vs. ps or p) is predominant.

#### The Case of Saturating Excitability

The saturable case is simpler than the non-saturable case. When pr + ps + 6p exceeds 1, all domains become active if pr > 0. However, a significant slowing down effect occurs when c is close to 1 − pr, so the time taken to activate all domains becomes long enough to make the seizure pattern appear non-global during a clinical observation period (Fig 5A).

### Other Consequences of Criticality

#### Pre-ictal Desynchronization

When increasing excitability pushes the system closer to its phase transition and thereby creates enlarging clusters, we speculate that the normal background synchrony becomes disrupted even when cluster diameters are small. Although the domains within a particular cluster may be synchronized, global synchrony in this case is low because there is little synchrony between the disconnected clusters, and the EEG thus exhibits a pre-ictal desynchronization. As the system draws closer to its phase transition, the clusters enlarge and coalesce, so that their domains are able to synchronize across a wide expanse of cortex, producing high amplitude EEG oscillations. Others have also advanced this hypothesis [25, 59].

#### Sudden Onset and Cessation of Seizures

Sudden onset or cessation of seizures may be due to several mechanisms, including those that increase excitability through increases in ps or p. These changes push the system closer to criticality, thereby inducing a high sensitivity to physiological parameters. Fig 3D shows examples in which m(t) decreases suddenly and significantly in response to small decreases in pr or p, and in which a small increase in pr can transform a grid with little activity to one with full activation. Although the percentage change of pr and p in these examples is high, the absolute changes are quite small, and could be produced by small or random changes in physiological variables.

### Effects of Discrete Foci

Seizure foci were modeled either as single domains or as 10 × 10 regions activated for a number of time steps. In the non-saturating case, a transient seizure focus has little effect on the final percentage of active domains, because Equation 1 applies after the focus vanishes. For example, all domains must eventually deactivate if pr = 0. However, the route to global deactivation can be very long and complex, due to slowing down effects if the system is near criticality (Fig 5B). This is a possible explanation of how a small, transient focus can produce significant and lengthy seizure activity prior to complete deactivation.

In the case of saturating excitability, the simulated focus led to global activation even if pr = 0. Again, slowing down effects can render the route to full activation both long and complex (Fig 5A). In this case of high excitability, a small, transient focus produced a slow route to complete activation.

### Addition of Refractory Periods

In other simulations, domains entered a *refractory period* after activation for longer than a *refractory threshold*. Typical refractory periods and refractory thresholds were 30 and 10 time steps, respectively.

#### Transitions from Clusters to Waves

When activity of a saturating system was induced by a transient stimulus rather than spontaneous activation (i.e., when pr = 0), and when ps and p were relatively low, refractory periods were absent because the duration of domain activity was less than the refractory threshold. The result was an expanding mass of active domains (Fig 6A). As p increased, longer durations of activity triggered refractory periods that disrupted this expanding mass and prevented full activation. In this case we observed slowly drifting clusters similar to ‘blobs’ reported in models of gap junctions (Fig 7) [38]. These movements were due to 3-dimensional clusters arising within the equivalent 3-dimensional Ising model that appeared as a sequence of clusters at successive times in our 2-dimensional model (see Methods and [47]), and may be the basis of the disorganized wave propagation observed in human recordings [60]. In contrast, the addition of refractory periods to non-saturating models did not lead to well-formed waves.

**Fig 6.**
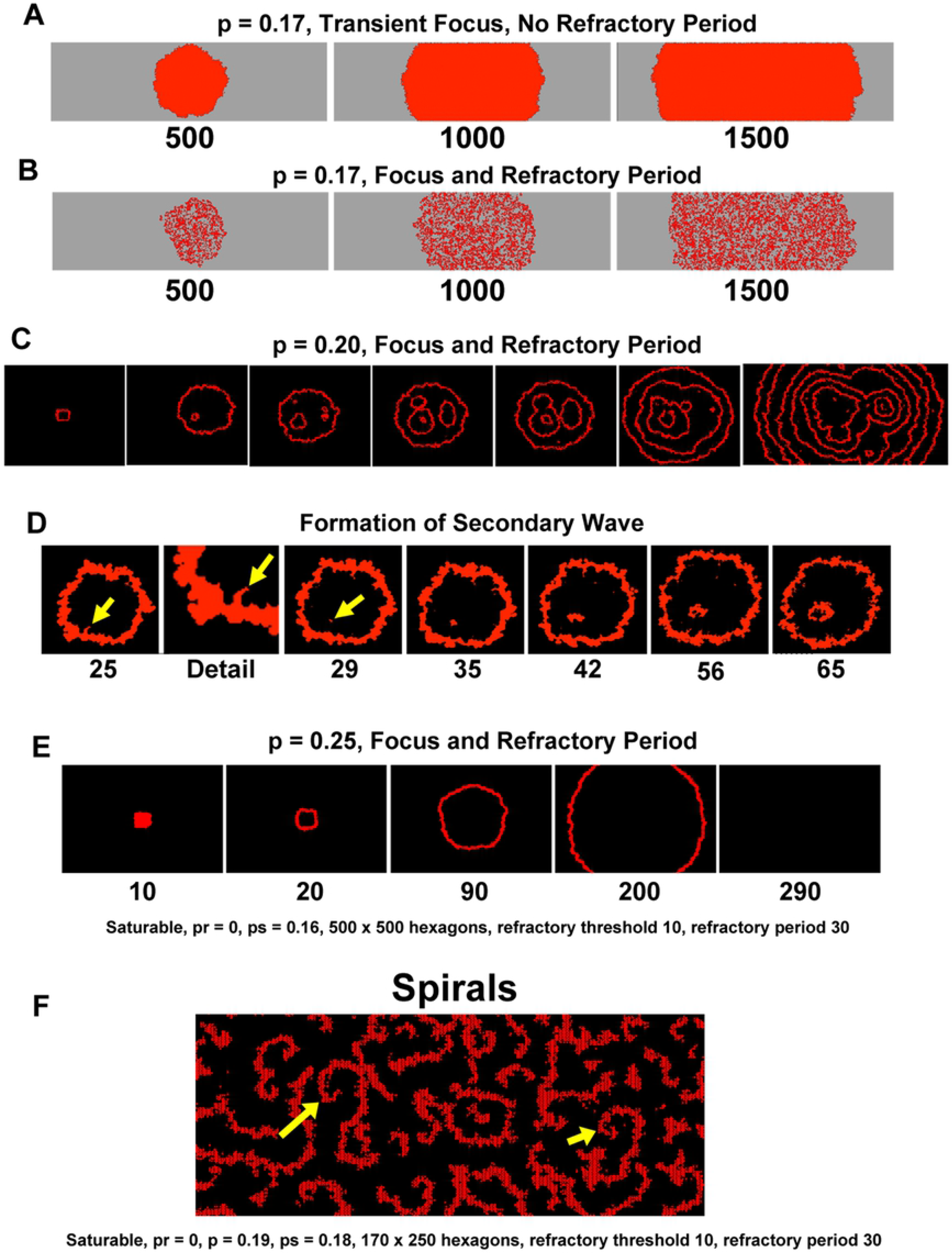
Refractory Periods Produce Waves and Secondary Waves. (A) Example of how a transient focus produces a simple growth pattern in saturating systems. Numbers indicate time step. (B) Same as (A), with refractory period added. Growth pattern persists but is disrupted by synchronizing effect of refractory period. (C) Same as (B), with p increased. Synchrony is high enough to produce persistent, well-formed waves. Note the presence of new secondary waves. (D) Detail to show how secondary waves are spawned. A small outgrowth from the primary wave (yellow arrow, 25 and Detail) becomes isolated (29), and grows to a cluster that forms a new ring due to a refractory center. These events do not always occur due to the statistical nature of the spread. (E) When p is increased further, secondary waves do not occur (see text). (F) Simulation producing spiral patterns (arrows).

**Fig 7.**
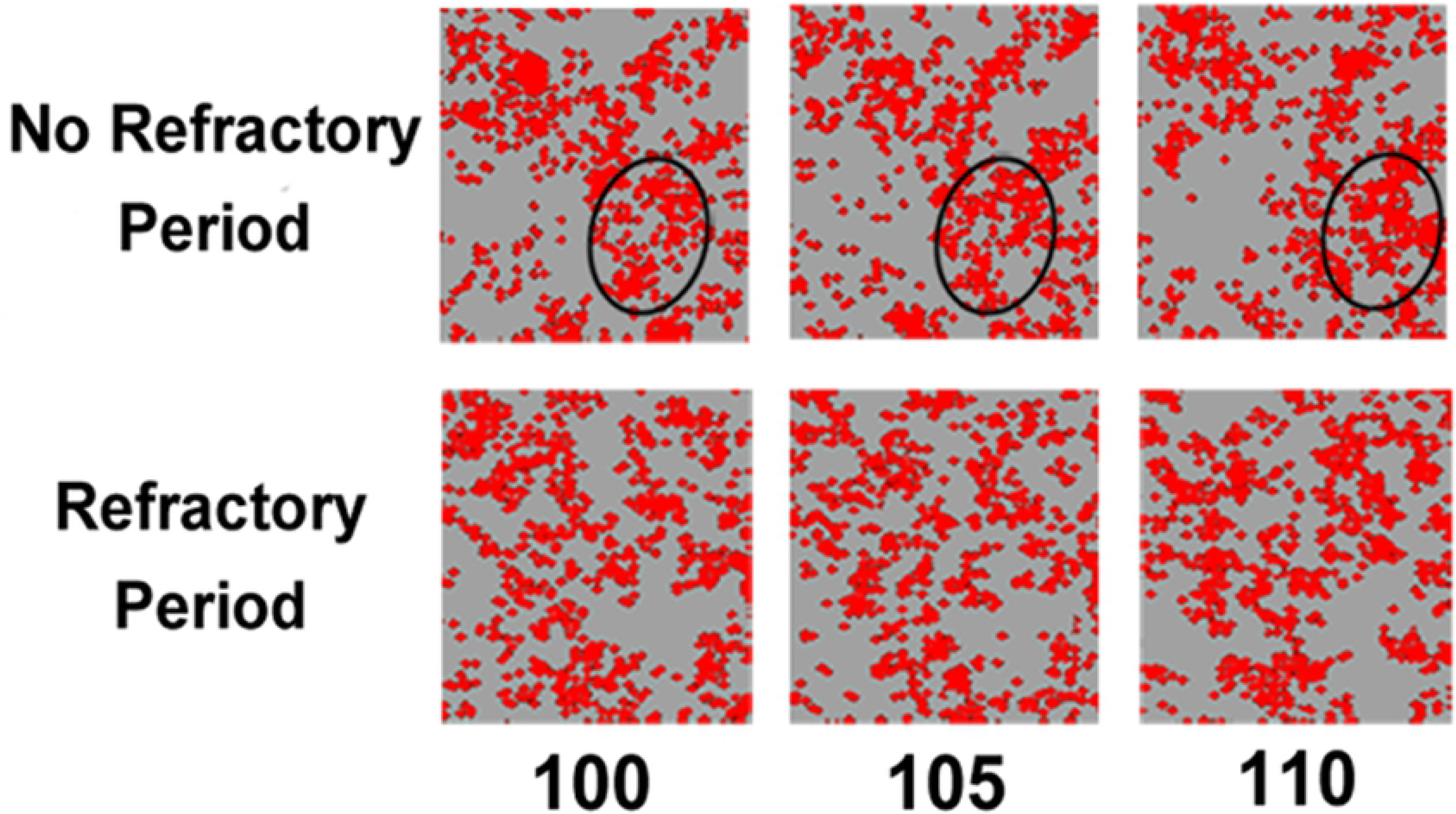
Cluster Drift. Top row shows cluster (in oval) drifting slightly as time increases from 100 to 110 steps (no refractory periods). No drift is seen when refractory periods are present (bottom row).

Further increases in p transformed the drifting clusters into poorly formed waves, and then into well-formed waves and spirals (Fig 6). The refractory periods seemed to promote synchronization by inhibiting the global activation would otherwise have occurred. When p was very high, the complexity vanished to produce a single, concentric wave vanishing at the boundary (Fig 6E), modeling the case of a single, transient seizure focus within an excitable neocortex in the presence of refractory periods.

The waves produced by our model were similar to those of other models and to those observed *in vivo* [1, 3, 4, 6, 26, 28, 36, 38, 40, 61, 62], including plane waves, concentric waves emanating from a focus, secondary waves spawned from wave collision or from the patch boundary, annihilation of colliding waves, and spiral patterns. The gradual transition between clusters and waves may explain reports of microseizure activity fluctuating between clusters and ‘expanding and contracting’ waves with variable degrees of organization [18].

It seems paradoxical that increases of p can change a pattern of complex, ongoing waves into a single concentric wave, i.e., that increases in excitability can lead to improvement of the seizure pattern. This occurred because high values of p simultaneously activate domains along the trailing edge of the wave, producing a smooth arc of activation followed by a smooth arc of refractory domains, which made secondary waves unlikely. For lower values of p, the edges were irregular and allowed the formation of secondary waves (Fig 8). These observations provide an explanation for the observation that increases of inhibitory drive can lead to a more severe and prolonged seizures [63].

**Fig 8.**
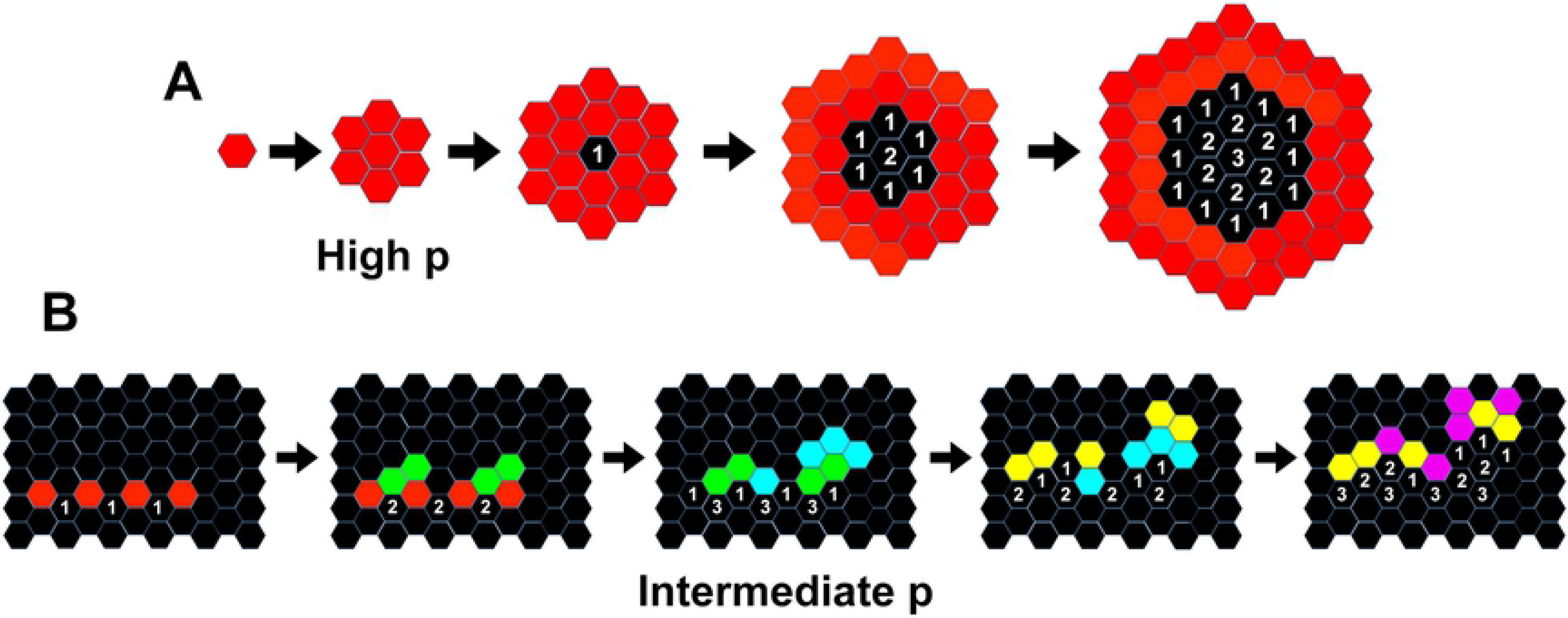
Effects of Excitability on Wave Edges. (A) Diagram to show how high values of p produce waves without secondary waves. Because p is high, all surrounding domains become active at each time step. The inner domains become refractory, creating a core of deactivated domains. If pr = 0, the domains in the core cannot reactivate and an expanding wave is produced. Numbers indicate the time stage of the refractory period. (B) Diagram to show how lower values of p can produce secondary waves. In this example, the refractory threshold is 2 and the refractory period is 3. Because p is not high, only a few domains activate around the active domains, producing an irregular wave front. Clusters occur that become isolated from each other due to the transformation of active domains into refractory domains. Each isolated group can spawn secondary waves. Red indicates active domains at the beginning, with domains activating in four subsequent stages indicated by the colors green, blue, yellow and purple.

When spontaneous activation was allowed (i.e., when pr > 0), drifting clusters were again observed for low values of p. As p increased the entire grid ‘flickered’ between a global activation with faint wave patterns and an almost complete absence of activation. Whether this pattern exists *in vivo* is uncertain.

Not all choices of refractory periods produced waves, in agreement with other studies [6]. Our observation of the transition to waves echoes the Hopf bifurcations produced in models of neural fields with refractory periods [64–66].

### Self-Perpetuating Waves

In a pioneering model of atrial fibrillation, Wiener and Rosenblueth showed that a small “dead zone” of no activity representing an infarct could induce circular, reentrant waves within an excitable myocardium [67]. To raise the possibility of similar behavior within the neocortex, we followed their work by placing two foci between two dead zones, timed so that the refractory periods of the waves from the first focus inhibited one of the waves from the second focus. This produced a single, unpaired wave, travelling repeatedly through and around the dead zones (See Methods) (Fig 9). These results suggest self-perpetuating seizure waves may occur within regions harboring complex, microscopic arrangements of non-functioning neocortex, and provide another possible mechanism for EEG oscillations and recurring, macroscopic neocortical seizure waves [1, 4].

**Fig 9.**
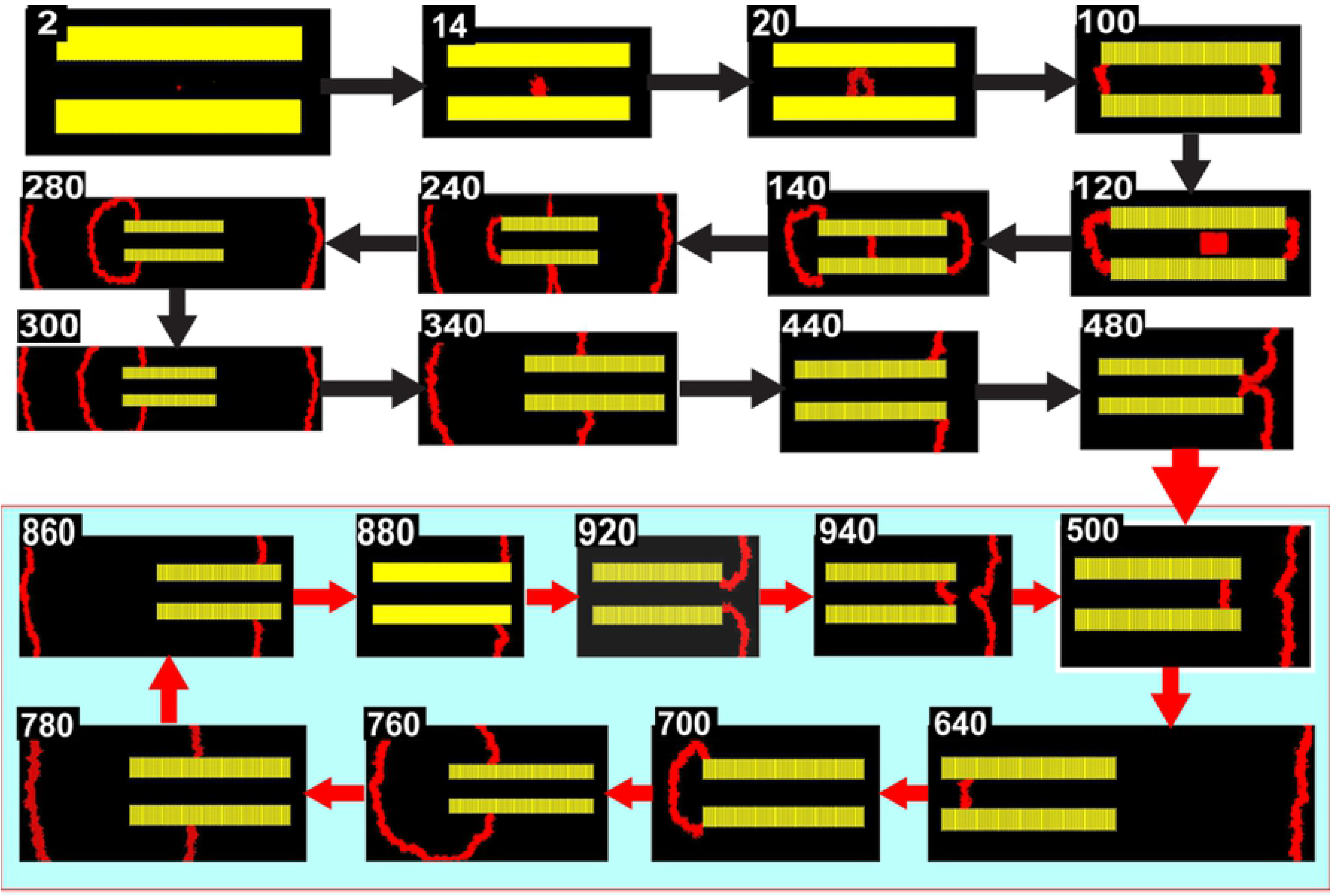
Recurrent Seizure Loops. Yellow bars are ‘dead zones’ in which domain activation is forbidden. Excitability is high and the refractory period is large (pr = 0, p = 0.25, ps = 0.15, c = 1.65, refractory threshold = 10, refractory period = 80). A small focus occurs for a single time step (step 2), and a second, larger focus occurs for a single time step later in the simulation (step 120). The initial focus grows and splits into two waves traveling along the dead zone in different directions (steps 14 to 140). Each wave splits three components. One component leaves the grid, and the other two annihilate their counterparts from the other wave (step 240 to 300). The second focus generates a single wave rather than two waves because of the refractory domains produced by the waves from the first focus. This wave travels between the dead zones (step 140), curling at the end to split into three waves (step 280 to 340). One of the waves leaves the grid (step 340 to 440), and the other two are not annihilated as before because the only one wave appears in step 140. The two waves instead rejoin at the opposite end to form the pattern seen at step 500. After step 500, the component between the dead zones cycles repeatedly, producing two waves in opposite directions during each cycle (the repeating cycle is shown with a blue background). Because of the statistical nature of the simulation, this behavior is not seen with every run.

## Discussion

A major assumption of our model is that probabilities can be assigned to the occurrence of seizure spread between adjacent domains, to the spontaneous creation of a seizure within a domain, and to the persistence of a seizure within a domain between time steps. These are simple rules, but lead to complex dynamics that require methods borrowed from statistical physics and the theory of criticality for analysis. We used these methods to show that the model mimics a surprisingly wide spectrum of seizure behaviors that would be difficult to otherwise subsume under a single theory. These behaviors occur at both microscopic and macroscopic scales, and include the high sensitivity of some seizures to small physiological changes, complex responses to transient stimulation, the occurrence of sudden seizure onset and cessation, the long time required for some seizures to reach their peak effects, the presence of enlarging clusters of microseizures, preictal desynchronization, the presence of self-limiting and recurrent seizure waves, the presence of drifting ‘blobs’, and the sometimes observed paradoxical worsening of seizures with increased inhibition. The model links microscopic dynamics with macroscopic events, thereby offering a way to reconcile the seemingly contradictory view of seizures as discrete clusters of hyperactive neurons vs. waves, and also yielding a possible explanation for how macroscopic seizure waves emerge from microscopic dynamics. These findings support our hypothesis that the theories of statistical physics and criticality can provide insight into the origins of macroscopic seizure behavior.

The other assumptions of our model, such as the presence of refractory periods, are also simply stated, allowing the model to be simply described with a small number of variables and rules with intuitive meaning. To our knowledge, few seizure models have produced such a wide variety of behavior from intuitive foundations, or without the need for a large number of less intuitive variables or a large number of differential equations. However, we stress our model does not replace detailed or neural mass models, nor does it simulate neurophysiological recordings. Furthermore, our model is clearly not fully realistic for reasons that include its construction as a two-dimensional grid. Moreover, its approach is semi-qualitative, similar to others in which biophysical mechanisms are ‘mapped’ onto model parameters, as in the use of dynamical systems to elucidate the behavior of seizure onset and cessation [31, 32, 68, 69]. Nevertheless, that our model produces so many different types of seizure behavior at different spatial and temporal scales suggests that its rules and assumptions may at least partially describe the underlying biological mechanisms, and thus confers insight into seizure dynamics.

These insights include possibilities for seizure treatment. Elevations in pr, ps or p represent different types of excitability that might be exploited to design clinical intervention. For example, systems with small pr and m will have few active domains, and thus minimal signs of seizures. But if the system is near criticality, the resulting high sensitivity of the system allows seizures to be easily triggered from deceptively small variations within a benign resting state. In this case, reducing ps or p will reduce c, move the system away from criticality, and be more effective than reducing pr (point ‘a’ in Fig 3A). This state could be detected by the presence of large clusters or long recovery times to stimulation, consistent with reports that delayed responses are associated with neocortical excitability [70]. On the other hand, for systems far from criticality, a reduction in pr can be much more effective than reductions in ps or p (point ‘b’ in fig 3A). This situation could be detected by a diffuse pattern of activation with small clusters.

### Translating Criticality into Seizure Behavior

The sensitivity of our model to small changes mirrors the viewpoint that seizure transitions occur as small fluctuations move the system on or off an attractor of a dynamical system [31, 32, 66, 68]. For example, small changes in model probabilities can produce or destroy global activation if the system is near criticality. (Fig 3D). Sensitivity to small changes due to phase transitions has also been found in local field potentials at seizure cessation, but not in microelectrode recordings (26). Because mean field estimates of criticality can be inaccurate [45], it is possible that phase transitions do not occur at the microseizure level, in agreement with our findings. Our observation of ‘flickering’ between high and low activation is a related hallmark of criticality, occurring *in vivo* and in other seizure models [28, 33, 35].

Other aspects of criticality also have clinical consequences. ‘Slowing down’ produces longer durations of activity, explaining how a small transient focus could produce long and complex seizures that eventually vanish if m = 0, or end with global activation if m ≥ 1. Slowing down also occurs in other models of seizure cessation [24, 28, 31]. Increases of correlation length and formation of clusters associated with criticality provide an explanation for the presence and coalescence of clusters of microseizures [11–15, 22–24]. Others have noted this process may explain preictal desynchronization: clusters that are at first small, disconnected and asynchronous yield a low amplitude EEG, but yield typical EEG recordings of higher amplitude as they coalesce and synchronize [24, 26].

### Clusters, Waves and Reentrant Behavior

Refractory periods triggered a transition between sparsely distributed clusters and seizure waves, in agreement with reports of fluctuations between clusters and complex ‘collapsing or expanding’ waves [18]. We speculate that excitation of an ictal core [27] could create waves in the same fashion, while feed-forward inhibition could decrease the duration of domain activation below the refractory threshold, thus inhibiting refractory periods and waves in the ictal penumbra.

Our model produced waves (Fig 6) similar to those of computational and *in vivo* studies, including plane and concentric waves, waves spawned from wave collision or from the patch boundary, and colliding waves with annihilation. Furthermore, the prolonged duration of wave activation arising from decreases in excitation is consistent with the well-known paradoxical worsening of seizures associated with increased inhibition.

Following early models of atrial fibrillation [67], we produced reentrant waves traveling in circular patterns. Such patterns are thought to explain delayed responses to seizure foci as ‘loops’ of reentrant activity [70], and might arise from ‘regional anisotropies’ [4] such as those associated with cortical dysplasia.

### Rationale for Refractory Periods

We included refractory periods because refractory behavior is likely as metabolites become exhausted, and because ‘pockets of refractoriness’ produce wave-like progression of interictal spikes [71]. Furthermore, refractory periods have been implicated in the formation of seizure waves by mathematical modeling [66], and ‘ictal inhibitory restraint’ may play a role in the production of waves [11]. Other models also include refractory periods [35, 40, 71].

Waves occur in mathematical models due to refractory behavior alone [34], and seizure waves may occur in models solely from an underlying mathematics. However, refractory periods are not needed for wave production [72], although some continuous models producing waves without refractory periods are equivalent to discrete models with refractory periods [73].

### Arrangement of Domains

A discrete nature of microscopic seizure activity is supported by the observation of microseizures [12–15, 21], ictal calcium maps of submillimeter clusters of hyperactive neurons [11], the finding that spikes propagate through the entorhinal cortex in steps of 0.5 mm [74], and the occurrence of local neuronal connectivity in discrete submillimeter regions [75]. A discrete form of seizure spread is also included in several mathematical models [6, 37, 38]. Our use of discrete domains was based on these considerations and reflects a common view that seizures can spread along anatomic pathways defined by columnar organization.

These considerations and studies of the arrangement of cortical columns motivated our use of a hexagonal grid [48, 49, 76]. Although the true arrangement of columns is not likely to be perfectly hexagonal, our results do not change if triangles or rectangles are used instead of hexagons, and the production of waves from refractory periods is robust to lattice changes [77]. These arguments suggest that our results apply to grids of modest irregularity, but the exact degree of regularity needed for validity of the model requires further study.

#### Limitations

We modeled an isolated cortical patch at equilibrium, without external inputs, knowing that such a patch does not exist *in vivo*, but believing that understanding this case would be a necessary first step. Further development such as long-range connections will likely require techniques from non-equilibrium physics. We assumed that the model probabilities are determined by physiological parameters, much as how physiological parameters are thought to determine variables of dynamical systems used to model seizure behavior [68]. This is supported by observations of probabilistic propagation at the microscopic level [78]. However, these assumptions and those of stability of model probabilities over short intervals need validation, perhaps using recordings of large numbers of microseizures to obtain probability estimates.

#### Conclusion

In a model combining probabilistic seizure spread, neocortical modularity, and the possibility of refractory periods, we used methods of statistical physics and criticality theory to demonstrate the presence of a continuous phase transition, and showed that the hallmark properties the phase transitions offer explanations for a wide variety of seizure behavior. The model offers possible explanations for the coexistence of seizure waves and clusters of activity, for the emergence of seizure waves from microscopic dynamics, and for a surprisingly wide variety of known properties of seizure. The model is not fully realistic, but the large number of complex seizure behaviors that it mimics suggest that similar mechanisms may occur *in vivo*. Because of its computational efficiency and use of a small number of intuitive parameters, the model is well suited for computational experimentation. Future work is needed to validate the model assumptions, but better understanding of how macroscopic seizure behavior emerges from microscopic seizure dynamics could inform both the understanding of seizure spread and the development of interventions such as the use of antiepileptic drugs that exert their effects at the microscopic scale to produce changes in macroscopic behavior.

## Acknowledgments

The author thanks David Blake and June Yowtak for insightful discussions and comments.

## Methods

### Calculation of Probabilities

In simulations, the state of each domain at each time step was calculated by first choosing a random number r for each domain and activating the domain if its associated r is less than its probability of activation. Domains were simultaneously updated at each time step. Simulations were performed with custom software in the Matlab environment. In most cases, thousands of steps could be achieved within a few minutes using a laptop computer.

### Calculation of m for non-saturating models without permanent foci

Here we show that for non-saturating models, the fraction m(t) of active domains at time t asymptotically approaches the value

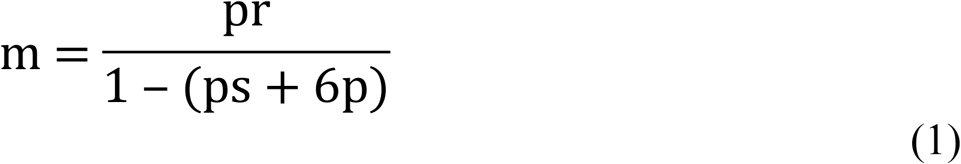

as t increases. In the absence of permanent seizure foci and refractory periods, the probability of a domain becoming active at time t depends on its state and those of its surrounding domains at time t − 1. This conditional probability is written as P(1|S), where S indicates the collective states of the domain and its neighbors at the previous time step. For the moment, we use values of 0 and 1 for domain states, with the value of 1 indicating an active domain.

If pr = 0, all domains remain inactive, m remains 0 and Equation 1 is valid. Note that any new activation such as a seizure focus will produce a robust activation if ps or p are high. We thus assume that pr is non-zero, and that pr + ps + 6p < 1 because the system is non-saturating. For reasons that will become clear, we now rewrite the system so that domain states take values of −1 and 1 (instead of 0 and 1), with 1 indicating activation. We will write s_i_ for domain states when using −1 and 1, reserving S_i_ for domains states when using of 0 and 1. Note that the index i ranges between 0 and 6. Using i = 0 refers to the state of the domain in question, and values of i between 1 and 6 refer to states of the surrounding domains. A PCA is said to be affine if there are constants V and T_i_ such that P(1|S) = ½(1 + V + ΣT_i_s_i_), where the sum ranges over the index i and s_i_ ranges over the states of the domain and its neighbors at time t − 1 [47, 56]. Note that this condition is equivalent to P(1|S) − P(−1|S) = P(1|S) = V + ΣT_i_s_i_ (using P(−1|S) = 1 − P(1|s)). We first show that our model is an affine PCA.

P(1|S) is the probability that a domain will be 1 given the state S at the prior time step. In our model, it is defined to be pr + psS_0_ + pΣS_i_, where S_0_ is the state (0 or 1) of the domain in question and S_i_ are the states of the six surrounding domains at time t − 1. To address the case in which domains assume values of −1 or 1, we define s_i_ = 2S_i_ − 1 and note that s_i_ = −1 (respectively, 1) when S = 0 (respectively, 1). Substituting, we find P(1|S) = pr + ps(s_i_ +1)/2 +pΣ(s_i_ +1)/2. To determine if this equation defines an affine system, we ask whether this expression can be written as ½(1 + V + ΣT_i_s_i_) for some values of V and T_i_.

Setting the variable terms equal, we obtain pss_i_/2 + pΣ s_i_/2 = ½(ΣT_i_s_i_), i.e., T_0_ = ps and T_i_ = p, for i ranging between 1 and 6. This shows that ΣT_i_ = ps + 6p. Setting the constant terms equal, we obtain pr + ps/2 + 6p/2 = ½(1 + V), so that V = 2pr + ps + 6p − 1 = 2pr + ΣT_i_ **−1**. Our model is therefore affine. Note also that the because pr > 0 and pr + ps + 6p ≤ 1, then ps + 6p = ΣT_i_ must be less than 1.

Now let H(t) be the average sum of states of the domains over the grid, i.e., the sum of the 1’s and −1’s at time t. For example, H(t) = −1 if all domains are inactive at time t. We seek to calculate m = H(∞) = the limit of H(t), as t grows large.

For an affine PCA, it can be shown that H(t+1) = V + H(t)ΣT_i_ [47, 56]. Iterating from H(0), we obtain

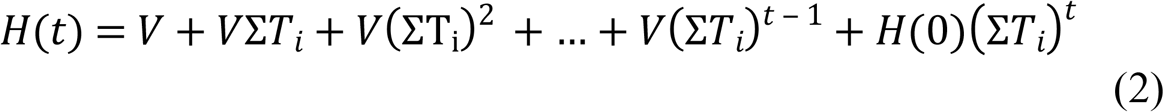

Because ΣT_i_ < 1, the last term of the right-hand side converges to 0 as t increases, and the remaining infinite series converges to H(∞) = V/(1− ΣT_i_).

H is the average value of the sum of the domain states when the states are either −1 or 1. Expressing H(∞) for the case where the states are 0 or 1 yields m, and is achieved by transforming H(∞) with the inverse of the formula used above to transform states of 0 and 1 to those of −1 and 1, i.e., m = (1 + H(∞))/2 = (1 + V/(1− ΣT_i_))/2 = pr/(1 − ΣT_i_). Using ΣT_i_ = ps + 6p and V = 2pr + ΣT_i_ − 1 derived above, we find m = pr/(1 − (ps + 6p)), as asserted in Equation (1).

It can also be shown that the explicit form for H(t) is H(t) = H(∞) (1 − (ΣT_i_)^t^) when the grid is initially non-active (see Slowing Down section below).

### Percentage of Active Domains for Saturating Excitability Without Refractory Periods

If pr = 0, m will be 0. However, for non-zero pr, all simulations of saturating systems (pr + ps + 6p ≥ 1) resulted in activation of every domain on the grid. We argue that this is intuitively plausible as follows. For any number m_0_ < 1, probabilities qr < pr, qs < ps and q < p can be found so that qr + qs + 6p < 1, and thus Equation 1 shows that the percentage of active domains of the non-saturating system represented by qr, qs and q is m0 = qr/(1-(qs + 6q)). Because the probability of domain activation at each step of the system represented by the p’s is greater than or equal to that of the system represented by the q’s, the percentage of active domains for the former system must be greater than m_0_ for any choice of m_0_ and hence must be at least 1.

### Slowing down

The arguments above show that H(∞) = V(1 + ΣTi + (ΣTi)^2^+…), and so

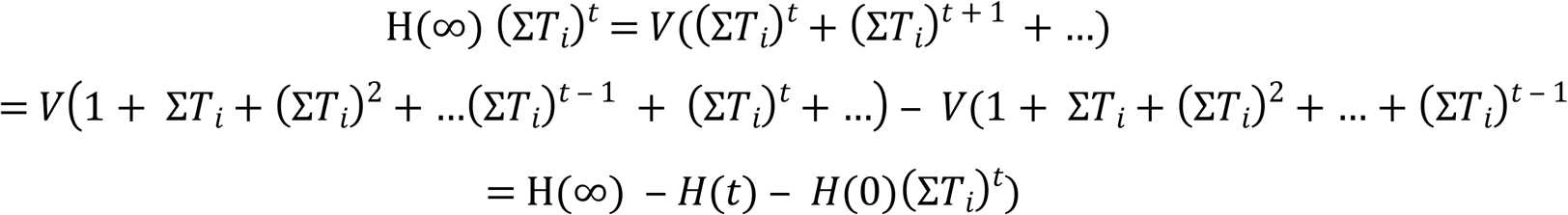

where the expression H(t) from the previous sections was used to derive the last step.

Collecting terms, we find that

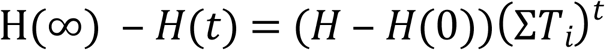

and so H(t) approaches H(∞) exponentially as confirmed in our examples. The half-life of this process occurs when (ΣTi)^t^ = ½, so the half-life t_1/2_ = −log(2)/log(ΣTi) = −log(2)/log(ps +6p). Therefore, when pr is small and as c = ps + 6p increases to its critical value of 1 − pr, the half-time of the system increases dramatically (Fig 3B and 3C), i.e., there is a ‘slowing down’ effect for H(∞). Because m = ½(1 + H(∞)), the is true for m.

For the saturable case, simulations showed that the time taken to activate all domains rapidly decreases as c increases beyond 1 − pr. The result was a slowing down as c approaches 1 − pr from either direction. (Fig 3B and 3C).

### Calculation of correlation lengths

Fluctuations of the seizure states of adjacent domains are often correlated due to local interactions, but correlations decrease exponentially as the distance between domains increases [43, 45]. The decay constant of this decrease is the *correlation length*. As a continuous phase transition is approached, the correlation length grows without limit, with high correlations found at all distances if the grid is infinitely large [30, 43, 45]. Because cortical patches are finite, correlations near critical points display finite peaks rather than infinite spikes. These peaks are strong evidence for a continuous phase transition.

To our knowledge, the correlation length of our model cannot be directly calculated (see Baxter [51] for a discussion of special cases admitting exact solutions). We therefore computed correlation lengths for several examples as follows [57].

In order to minimize the effects of the grid boundary, calculations were limited to a 120×120 block of domains centered within a 200×200 grid or a 300×300 region centered within a 500×500 grid. Furthermore, correlation calculations did not begin until the system had achieved its stable value of m, typically after 2000 to 3000 time steps. At each time step, for each domain c within the block, the domains at a distance r in each of the vertical and horizontal directions were labeled c1, c2, c3 and c4. We calculated the average of the four correlations (state of c − mt)(state of ci − mt), where mt was the average number of active domains at time t [57], and varied r between 1 and 60 (for 200×200 grids) or between 1 and 100 (for 500×500 grids) to produce a graph of the resulting correlations vs. r with an exponential shape. The smallest value of r yielding a correlation less than 0.001 was taken as the *correlation length* associated to the domain c. These values were averaged over all domains within the block to obtain the correlation length at that particular time step. This calculation was repeated for each time step until the correlation distances were stable, typically requiring 5000 to 10000 time steps, and the average value over the final 2000 time steps reported as the final correlation length.

Graphs of the correlation distances displayed peaks were found when c approached 1 − pr (Fig 4A, 4B and 4C), supporting the presence of a continuous phase transition.

The large correlation lengths near a continuous phase transition strengthen the long-distance correlations, thereby producing large clusters of correlated domains [45]. The appearance of such clusters in our model suggests that the clusters of microseizures observed in the microgrid data may occur through proximity to a continuous phase transition associated with increased excitability.

To illustrate, we compared the behavior of clusters in two systems with the same m, in which one system is close to criticality. The ‘critical system’ was defined by the probabilities pr = 0.009, ps = 0.19, and p = 0.132. The fraction of active domains m was 0.5 by Eq. (1), and ps + 6p = .982 was close (within 0.9%) to the critical value for c = 1 − pr = 0.991. The ‘non-critical’ system was defined by pr = 0.22, ps = 0.05 and p = 0.085, yielding m = 0.5 as before. The value for ps + 6p = 0.56 was relatively far (within 28%) from the critical value for c of 0.78. Twenty simulations were performed for each system for at least 600 time steps until m was stable (within 10% of calculated m). The average cluster area (i.e., the average number of grid nodes in each cluster) and number of clusters was calculated for each run with a density based clustering algorithm [61], and the results averaged and compared with a non-paired 2-tailed t-test (p < .001) for each system.

The non-critical system produced significantly more clusters than the critical system (1246.9 ± 20.4 vs. 683 ± 21.2, p < 0.001, mean ± sd), and the average size of its clusters was significantly smaller (13.5 ± 0.28 vs. 23.2 ± 1.1, p < 0.001). Cluster diameters from the non-critical system were less than 200, while the size of 232 of the clusters from the critical system exceeded 200 (Fig 4D).

### Self-perpetuating Waves

Following [67], two foci were activated between two rectangular dead zones [FIG]. The first was a single domain at the center of the grid, activated for a single time step. In most cases this produced two linear waves traveling in opposite directions between the dead zones. If no further focus was activated, each wave split at the ends of the dead zones, with two components extending towards the grid boundary and the other two components circling the dead zone to annihilate each other to stop all activity. We then added a second focus composed of a 15×7 rectangle of domains extending from 85 to 100 in the horizontal direction and 103-110 in the vertical direction, activated at time step 110. In most cases, the second focus also produced self-limiting concentric waves around the dead zones. In some cases, however, the refractory region behind one of the trailing edges of these waves blocked one of the annihilating pairs of waves, producing a single self-perpetuating wave travelling through and around the dead zones, spawning two travelling waves towards the boundary at each circuit (Fig 8).

## References

1. Huang X, Xu W, Liang J, Takagaki K, Gao X, Wu JY. Spiral wave dynamics in neocortex. Neuron. 2010;68(5):978–990.

2. Pinto DJ, Patrick SL, Huang WC, Connors BW. Initiation, propagation and termination of epilpetiform activity in rodent neocortex *in vitro* invlve distinct mechanisms. Journal of Neuroscience. 2005;25:8131–8140.

3. Vanleer AC, Blanco JA, Wagenaar JB, Viventi J, Contreras D, Litt B. Millimeter-scale epileptiform spike propagation patterns and their relationship to seizures. J Neural Eng. 2016;13(2):026015.

4. Viventi J, Kim DH, Vigeland L, Frechette ES, Blanco JA, Kim YS, et al. Flexible, foldable, actively multiplexed, high-density electrode array for mapping brain activity in vivo. Nat Neurosci. 2011;14(12):1599–1605.

5. Kramer MA, Kirsch HE, Szeri AG. Pathological pattern formation and cortical propagation of epileptic seizures. Journal of the Royal Society Interface. 2005;2:113–127.

6. Goodfellow M, Taylor PN, Wang Y, Garry DJ, Baier G. Modelling the role of tissue heterogeneity in epileptic rhythms. Eur J Neurosci. 2012;36(2):2178–2187.

7. Matsumoto H, Ajmone-Marsan C. Cortical cellular phenomena in experimental epilepsy: ictal manifestations. Experimental Neurology. 1964;9:305–326.

8. Babb TL, Wilson CL, Isokawa-Akesson M. Firing patterns of human limbic neurons during stereoencephalography (SEEG) and clinical temporal lobe seizures. Electroencephalography andl Clinical Neurophysiology. 1987;66:467–482.

9. Wyler AR, Ojemann GA, Ward AAJ. Neurons in human epileptic cortex: correlation between unit and EEG activity. Annals of Neurology. 1982;11:301–308.

10. Wyler AR, Ward AAJ. Neuronal firing patterns from epileptogenic foci fo monkey and human. Advances in Neurology. 1986;44:967–989.

11. Trevelyan AJ, Sussillo D, Watson BO, Yuste R. Modular propagation of epileptiform activity: evidence for an inhibitory veto in neocortex. J Neurosci. 2006;26(48):12447–12455.

12. Schevon CA, Ng SK, Cappell J, Goodman RR, McKhann G, Jr., Waziri A, et al. Microphysiology of epileptiform activity in human neocortex. J Clin Neurophysiol. 2008;25(6):321–330.

13. Schevon CA, Goodman RR, McKhann G, Jr., Emerson RG. Propagation of epileptiform activity on a submillimeter scale. J Clin Neurophysiol. 2010;27(6):406–411.

14. Stead M, Bower M, Brinkmann BH, Lee K, Marsh WR, Meyer FB, et al. Microseizures and the spatiotemporal scales of human partial epilepsy. Brain. 2010;133(9):2789–2797.

15. Truccolo W, Donoghue JA, Hochberg LR, Eskandar EN, Madsen JR, Anderson WS, et al. Single-neuron dynamics in human focal epilepsy. Nat Neurosci. 2011;14(5):635–641.

16. Bower MR, Stead M, Meyer FB, Marsh WR, Worrell GA. Spatiotemporal neuronal correlates of seizure generation in focal epilepsy. Epilepsia. 2012;53(5):807–816.

17. Kramer MA, Eden UT, Kolaczyk ED, Zepeda R, Eskandar EN, Cash SS. Coalescence and fragmentation of cortical networks during focal seizures. J Neurosci. 2010;30(30):10076–10085.

18. Wagner FB, Eskandar EN, Cosgrove GR, Madsen JR, Blum AS, Potter NS, et al. Microscale spatiotemporal dynamics during neocortical propagation of human focal seizures. Neuroimage. 2015;122:114–130.

19. Bragin A, Engel J, Jr., Wilson CL, Fried I, Buzsaki G. High-frequency oscillations in human brain. Hippocampus. 1999;9:137–142.

20. Bragin A, Mody I, Wilson CL, Engel J, Jr. Local generation of fast ripples in epileptic brain. Journal of Neuroscience. 2002;22:2012–2021.

21. Schevon CA, Trevelyan AJ, Schroeder CE, Goodman RR, McKhann G, Jr., Emerson RG. Spatial characterization of interictal high frequency oscillations in epileptic neocortex. Brain. 2009;132(Pt 11):3047–3059.

22. Bragin A, C.L. W, Engel J, Jr. Chronic epileptogenesis requires development of a network of pathologically interconnected neuron clusters: a hypothesis. Epilepsia. 2000;41:S144–S152.

23. Bikson M, Fox JE, Jefferys JG. Neuronal aggregate formation underlies spatiotemporal dynamics of nonsynaptic seizure initiation. J Neurophysiol. 2003;89(4):2330–2333.

24. Jiruska P, Csicsvari J, Powell AD, Fox JE, Chang WC, Vreugdenhil M, et al. High-frequency network activity, global increase in neuronal activity, and synchrony expansion precede epileptic seizures in vitro. J Neurosci. 2010;30(16):5690–5701.

25. Jiruska P, de Curtis M, Jefferys JG, Schevon CA, Schiff SJ, Schindler K. Synchronization and desynchronization in epilepsy: controversies and hypotheses. J Physiol. 2013;591(4):787–797.

26. Wang Y, Trevelyan AJ, Valentin A, Alarcon G, Taylor PN, Kaiser M. Mechanisms underlying different onset patterns of focal seizures. PLoS Comput Biol. 2017;13(5):e1005475.

27. Schevon CA, Weiss SA, McKhann G, Jr., Goodman RR, Yuste R, Emerson RG, et al. Evidence of an inhibitory restraint of seizure activity in humans. Nat Commun. 2012;3:1060.

28. Martinet LE, Fiddyment G, Madsen JR, Eskandar EN, Truccolo W, Eden UT, et al. Human seizures couple across spatial scales through travelling wave dynamics. Nat Commun. 2017;8:14896.

29. de Curtis M, Avoli M. GABAergic networks jump-start focal seizures. Epilepsia. 2016;57(5):679–687.

30. Beggs JM, Timme N. Being critical of criticality in the brain. Front Physiol. 2012;3:163.

31. Jirsa VK, Stacey WC, Quilichini PP, Ivanov AI, Bernard C. On the nature of seizure dynamics. Brain. 2014;137(Pt 8):2210–2230.

32. Kramer MA, Truccolo W, Eden UT, Lepage KQ, Hochberg LR, Eskandar EN, et al. Human seizures self-terminate across spatial scales via a critical transition. Proc Natl Acad Sci U S A. 2012;109(51):21116–21121.

33. Scheffer M, Carpenter SR, Lenton TM, Bascompte J, Brock W, Dakos V, et al. Anticipating critical transitions. Science. 2012;338:344–348.

34. Wilson H, Cowan J. Excitatory and inhibitory interactions in localized populations of model neurons. Biophysics. 1972;12:1–23.

35. Goodfellow M, Schindler K, Baier G. Intermittent spike-wave dynamics in a heterogeneous, spatially extended neural mass model. Neuroimage. 2011;55(3):920–932.

36. Wang Y, Goodfellow M, Taylor PN, Baier G. Dynamic mechanisms of neocortical focal seizure onset. PLoS Comput Biol. 2014;10(8):e1003787.

37. Traub RD, Schmitz D, Jefferys JG, Draguhn A. High-frequency population oscillations are predicted to occur in hippocampal pyramidal neural networks interconnected by ax-axonal gap junctions. Neuroscience. 1999;92:407–426.

38. Traub RD, Duncan R, Russell AJ, Baldeweg T, Tu Y, Cunningham MO, et al. Spatiotemporal patterns of electrocorticographic very fast oscillations (> 80 Hz) consistent with a network model based on electrical coupling between principal neurons. Epilepsia. 2010;51(8):1587–1597.

39. Wilson MT, Sleigh JW, Steyn-Ross DA, Steyn-Ross ML. General anesthetic-induced seizures can be explained by a mean-field model of cortical dynamics. Anesthesiology. 2006;104:588–593.

40. Goodfellow M, Schindler K, Baier G. Self-organised transients in a neural mass model of epileptogenic tissue dynamics. Neuroimage. 2012;59(3):2644–2660.

41. Friston KJ. On the modeling of seizure dynamics. Brain. 2014;137:2110–2118.

42. Proix T, Bartolomei F, Guye M, Jirsa VK. Individual brain structure and modelling predict seizure propagation. Brain. 2017;140(3):641–654.

43. Stanley HE. Introduction to Phase Transitions and Critical Phenomenon. New York: Oxford University Press; 1987.

44. Plishke M, Bergersen B. Equilibrium Statistical Physics. New Jersey: World Scientific; 2006.

45. Sethna JP. Entropy, Order Parameters and Complexity. Oxford: Oxford University Press; 2012.

46. Enting IG. Crystal growth models and Ising models: disorder points. Journal of Physics C. 1977:1379–1388.

47. Georges A, Le Doussal P. From equilibrium spin models to probabilistic cellular automata. Journal of Statistical Physics. 1989;54(3/4):1011–1064.

48. Gabbott PL. Radial organisation of neurons and dendrites in human cortical areas 25, 32, and 32’. Brain Res. 2003;992(2):298–304.

49. Muir DR, Da Costa NM, Girardin CC, Naaman S, Omer DB, Ruesch E, et al. Embedding of cortical representations by the superficial patch system. Cereb Cortex. 2011;21(10):2244–2260.

50. Ermentrout GB, Edelstein-Keshet E. Cellular automata approaches to biological modeling. Journal of Theoretical Biology. 1993;160:97–133.

51. Baxter RJ. Exactly Solved Models in Statistical Mechanics. New York: Dover; 2007.

52. Scheffer M, Bascompte J, Brock WA, Brovkin V, Carpenter SR, Dakos V, et al. Early-warning signals for critical transitions. Nature. 2009;461(7260):53–59.

53. Veraart AJ, Faassen EJ, Dakos V, van Nes EH, Lurling M, Scheffer M. Recovery rates reflect distance to a tipping point in a living system. Nature. 2011;481(7381):357–359.

54. Truccolo W, Ahmed OJ, Harrison MT, Eskandar EN, Cosgrove GR, Madsen JR, et al. Neuronal ensemble synchrony during human focal seizures. Journal of Neuroscience. 2014;34:9927–9944.

55. Meisel C, Klaus A, Kuehn C, Plenz D. Critical slowing down governs the transition to neuron spiking. PLoS Comput Biol. 2015;11(2):e1004097.

56. Rujin P. Cellular automata and statistical mechanical models. Journal of Statistical Physics. 1987;49:139–222.

57. Nagy TF, Mahant iSD, Tsallis C. Correlation function studies on the Domany-Kinzel cellular automaton. Physica A. 1998;250:345–354.

58. Domany E, Kinzel W. Equivalence of cellular automata to Ising models and directed percolation. Physical Review Letters. 1984;53:311–314.

59. Mormann F, Kreuz T, Andrzejak RG, David P, Lehnertz K, Elger CE. Epileptic seizures are preceded by a decrease in synchronization. Epilepsy Research. 2003;53(3):173–185.

60. Martinet LE, Ahmed OJ, Lepage KQ, Cash SS, Kramer MA. Slow Spatial Recruitment of Neocortex during Secondarily Generalized Seizures and Its Relation to Surgical Outcome. J Neurosci. 2015;35(25):9477–9490.

61. Steyn-Ross ML, Steyn-Ross DA, Sleigh JW. Gap junctions modulate seizures in a mean-field model of general anesthesia for the cortex. Cogn Neurodyn. 2012;6(3):215–225.

62. Bai L, Huang X, Yang Q, Wu JY. Spatiotemporal patterns of an evoked network oscillation in neocortical slices: coupled local oscillators. J Neurophysiol. 2006;96(5):2528–2538.

63. Paz JT, Huguenard JR. Microcircuits and their interactions in epilepsy: is the focus out of focus? Nat Neurosci. 2015;18(3):351–359.

64. Curtu R, Ermentrout B. Oscillations in a refractory neural net. Journal of Mathematical Biology. 2001;43(1):81–100.

65. Coombes S. Waves, bumps, and patterns in neural field theories. Biol Cybern. 2005;93(2):91–108.

66. Meijer HG, Coombes S. Travelling waves in a neural field model with refractoriness. J Math Biol. 2014;68(5):1249–1268.

67. Wiener N, Rosenblueth A. The mathematical formulation of hte problem of conduction of impulses in a network of connected excitable elements, specifically in cardiac muscle. Archivos del Instituto de Cardiologio de Mexico. 1946;16:205–265.

68. da Silva FL, Blanes W, Kalitzin SN, Parra J, Suffczynski P, Velis DN. Epilepsies as dynamical diseases of brain systems: basic models of the transition between normal and epileptic activity. Epilepsia. 2003;44:72–83.

69. Kramer MA, Szeri AJ, Sleigh JW, Kirsch HE. Mechanisms of seizure propagation in a cortical model. J Comput Neurosci. 2007;22(1):63–80.

70. Valentin A, Anderson M, Alarcon G, Garcia Seoane JJ, Selway R, Binnie CD, et al. Responses to single pulse electrical stimulation identify epileptogenesis in the human brain *in vivo*. Brain. 2002;125:1709–1718.

71. Sabolek HR, Swiercz WB, Lillis KP, Cash SS, Huberfeld G, Zhao G, et al. A candidate mechanism underlying the variance of interictal spike propagation. J Neurosci. 2012;32(9):3009–3021.

72. Ermentrout GB, Kleinfeld D. Traveling electrical waves in cortex: insights from phase dynamics and speculation on a computational role. Neuron. 2001;29:33–44.

73. Terman D, Ahn S, Wang X, Just W. Reducing Neuronal Networks to Discrete Dynamics. Physica D. 2008;237(3):324–338.

74. Stewart C. Columnar activity supports propagation of population bursts in slices of rat entorhinal cortex. Brain Research. 1999;830:274–284.

75. Voges N, Schuz A, Aertsen A, Rotter S. A modeler’s view on the spatial structure of intrinsic horizontal connectivity in the neocortex. Prog Neurobiol. 2010;92(3):277–292.

76. Cruz L, Urbanc B, Inglis A, Rosene DL, Stanley HE. Generating a model of the three-dimensional spatial distribution of neurons using density maps. Neuroimage. 2008;40(3):1105–1115.

77. Berry H, Fates N. Robustness of the critical behavior in the stochastic Greenberg-Hastings cellular automaton model. International Journal of Unconventional Computing. 2011;0:1–21.

78. Wenzel M, Hamm JP, Peterka DS, Yuste R. Reliable and Elastic Propagation of Cortical Seizures In Vivo. Cell Rep. 2017;19(13):2681–2693.

